# Robust fine-mapping in the presence of linkage disequilibrium mismatch

**DOI:** 10.1101/2024.10.29.620968

**Authors:** Wenmin Zhang, Tianyuan Lu, Robert Sladek, Josée Dupuis, Guillaume Lettre

## Abstract

Fine-mapping methods based on summary statistics from genome-wide association studies (GWAS) and linkage disequilibrium (LD) information are widely used to identify potential causal variants. However, LD mismatch between the external LD reference panel and the GWAS population is common and can lead to compromised accuracy of fine-mapping. We developed RSparsePro, a probabilistic graphical model with an efficient variational inference algorithm, to perform robust fine-mapping in the presence of LD mismatch. In simulation studies with a varying degree of LD mismatch, RSparsePro identified credible sets with a consistently higher power and coverage than SuSiE. In fine-mapping cis-protein quantitative trait loci, RSparsePro identified credible sets with a consistently higher enrichment of variants with functional impacts and cross-study replication rates. In fine-mapping risk loci for low-density lipoprotein cholesterol in ancestry-specific GWAS, RSparsePro identified biologically relevant variants in drug target genes and implicated potential regulatory mechanisms. RSparsePro is openly available at https://github.com/zhwm/RSparsePro_LD.

## 1 Introduction

Fine-mapping is a widely used statistical procedure for identifying potential causal variants for human traits and diseases. Several fine-mapping methods have been developed, utilizing the marginal variant-phenotype associations from genome-wide association studies (GWAS) and the correlation between variants (linkage disequilibrium, LD) to prioritize targets for functional follow-up studies [1, 2, 3, 4, 5, 6, 7, 8, 9, 10, 11]. For example, CAVIAR identifies potential causal variants by exhaustively evaluating the likelihoods of all possible causal configurations [4]. FINEMAP adopts a stochastic shotgun search algorithm that focuses on a subset of causal configurations, effectively reducing the computational cost [1]. SuSiE and SparsePro group correlated variants and jointly model their effects, significantly improving the efficiency and accuracy of fine-mapping [2, 3, 6].

Ideally, all these fine-mapping methods should utilize LD information derived from the same individuals as included in the GWAS population, which ensures accurate modeling of the correlation between genetic variants. However, due to ethical and logistical concerns, access to individual-level data is often limited, resulting in a lack of in-sample LD information. Consequently, fine-mapping typically relies on publicly available GWAS summary statistics and LD reference panels that consist of individuals from populations different from the GWAS population. Despite the intention to broaden the applicability of fine-mapping analyses, the use of external LD reference panels can lead to LD mismatch, where the LD structures differ between the LD reference panel and the GWAS population, even if the two populations share the same genetic ancestry [12].

In the presence of LD mismatch, existing fine-mapping methods can encounter convergence issues [2]. For instance, SuSiE utilizes LD information for both the prior estimation procedure and the iterative Bayesian stepwise selection algorithm [2]. Without closely matched GWAS summary statistics and LD information, both steps may fail to generate reasonable results [2]. Hypothesis testing-based quality control methods have been developed, such as DENTIST [12] and SLALOM [13], for detecting and excluding loci subject to LD mismatch. However, it is still desirable to have a robust fine-mapping method, which allows for potential LD mismatch and expands the applicability of fine-mapping analyses, especially with emerging large-scale GWAS that often include protected datasets or involve multiple cohorts and diverse populations.

In this study, we propose Robust SparsePro (RSparsePro) for fine-mapping in the presence of LD mismatch. In RSparsePro, we introduce a latent variable to account for inconsistencies between the LD reference panel and GWAS summary statistics and develop an efficient variational inference algorithm to enable robust fine-mapping. We evaluate the performance of RSparsePro in comparison with SuSiE through simulation studies with a varying degree of LD mismatch. Additionally, we illustrate the ability of RSparsePro to robustly identify biologically meaningful candidate causal variants by fine-mapping loci identified in GWAS for plasma protein levels [14, 15] and plasma low-density lipoprotein (LDL) cholesterol levels [16], where in-sample LD information is unavailable.

## 2 Results

### 2.1 RSparsePro for robust fine-mapping

The assumed data generating process in RSparsePro is illustrated in **Figure** 1. RSparsePro utilizes GWAS summary statistics and an LD reference panel as input. To perform robust fine-mapping accounting for potential discrepancy between the LD reference panel and the GWAS population, we introduce a latent variable in the RSparsePro model. Specifically, we assume that the observed z-scores in GWAS summary statistics 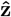 are error-contaminated observations of the latent z-scores **z**:

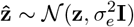

where 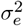 is a hyperparameter quantifying the magnitude of the discrepancy.

Next, similar to SuSiE [2, 6] and our previous work [3, 17, 18], we assume that within a locus containing *G* variants, there are up to *K* causal signals and the latent z-scores **z** are marginal variant-phenotype associations aggregating all of the *K* causal signals:

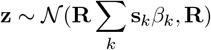

where **s**_*k*_ ∼ **Multinomial** 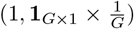 is the sparse indicator for the causal variant in the *k*^*th*^ credible set and 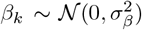 is the effect size of the *k*^*th*^ credible set.

To perform robust fine-mapping, our goal is to infer **s**_*k*_ given the observed z-scores 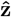 and the LD matrix **R** derived from the LD reference panel. We develop an efficient variational inference algorithm with closed-form expressions of iterative updates [19, 20] detailed in the **Supplementary Notes**. We use an uninformative prior for *β*_*k*_ to ensure that the prior estimation procedure remains unaffected by LD mismatch. The choice of other hyperparameters and the construction of 95% credible sets (used throughout this work) are detailed in the **Supplementary Notes**.

**Figure 1.**
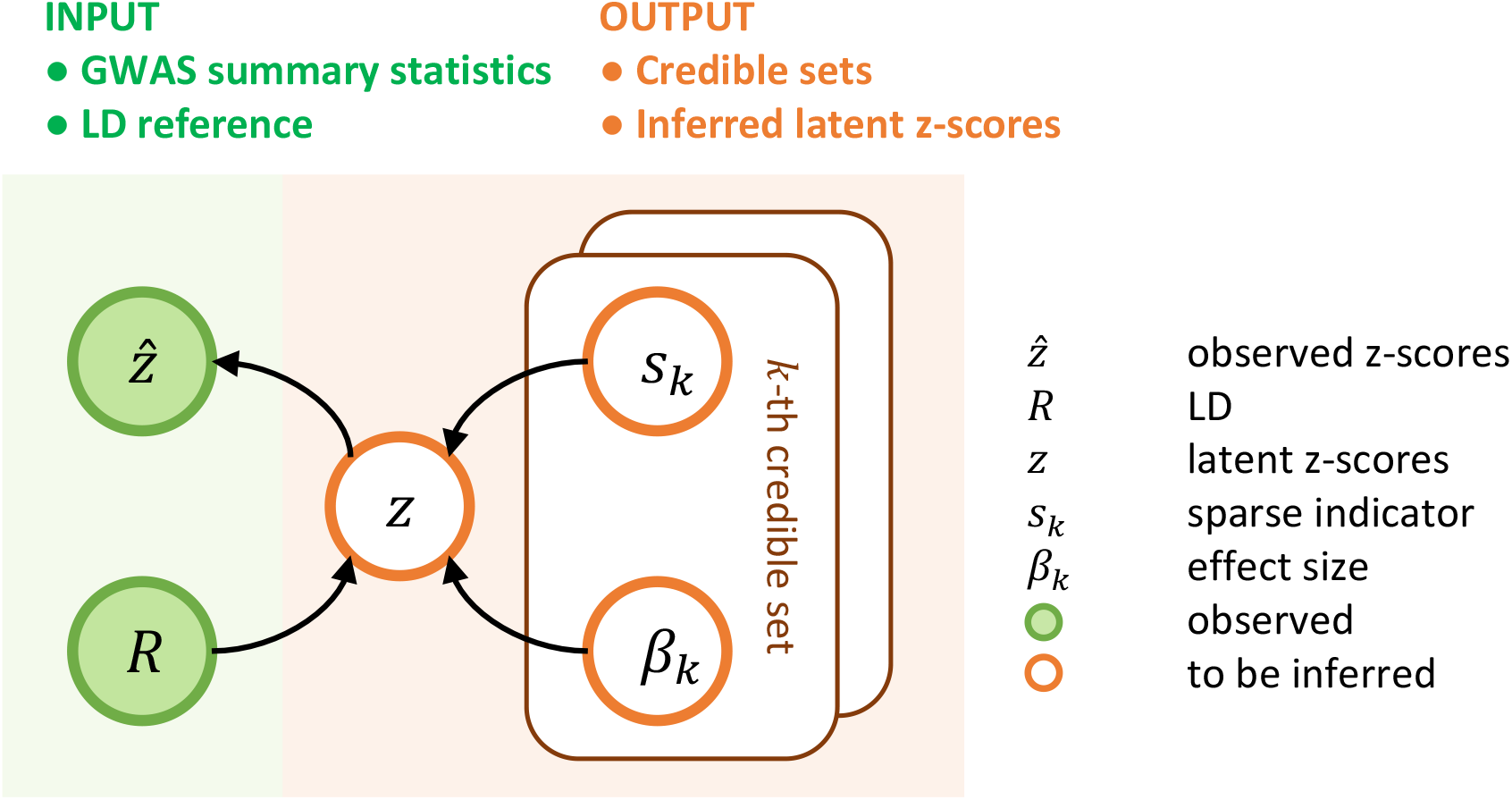
RSparsePro for robust fine-mapping. This graphical model depicts the data generating process in RSparsePro. Green shaded nodes represent observed variables, including the observed GWAS z-scores 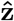 and the LD matrix **R** derived from a reference panel. Orange unshaded nodes represent variables to be inferred. **s**_*k*_ is a sparse indicator specifying the variant representation in the *k*^*th*^ credible set and *β*_*k*_ represents the standardized effect size of the *k*^*th*^ credible set. **z** is the latent z-scores. The variational inference algorithm for efficient and accurate posterior inference is detailed in the **Supplementary Notes**.

### 2.2 Robust fine-mapping in the presence of LD mismatch

To evaluate the performance of RSparsePro, we simulated GWAS z-scores both without and with a varying degree of LD mismatch (**Methods**). We categorized all simulation replicates based on whether SuSiE encountered convergence issues. SuSiE does not provide fine-mapping results when its prior estimation fails. One simulated example is presented in **Figure** 2A-C, with three causal variants and 10% of the variants having a mismatch factor of −0.5 (*K*_*c*_ = 3, *p* = 0.1, *f* = −0.5). Without LD mismatch (**Figure** 2A), both SuSiE and RSparsePro correctly identified all three causal variants in their respective credible sets. By multiplying the z-scores of 10% of the variants by a mismatch factor of −0.5 (**Figure** 2A and B), we introduced inconsistencies between the GWAS summary statistics and the LD reference panel. The prior estimation in SuSiE failed as a result of the simulated LD mismatch. In contrast, using the same input, RSparsePro correctly identified three credible sets containing the three causal variants, respectively (**Figure** 2C).

**Figure 2.**
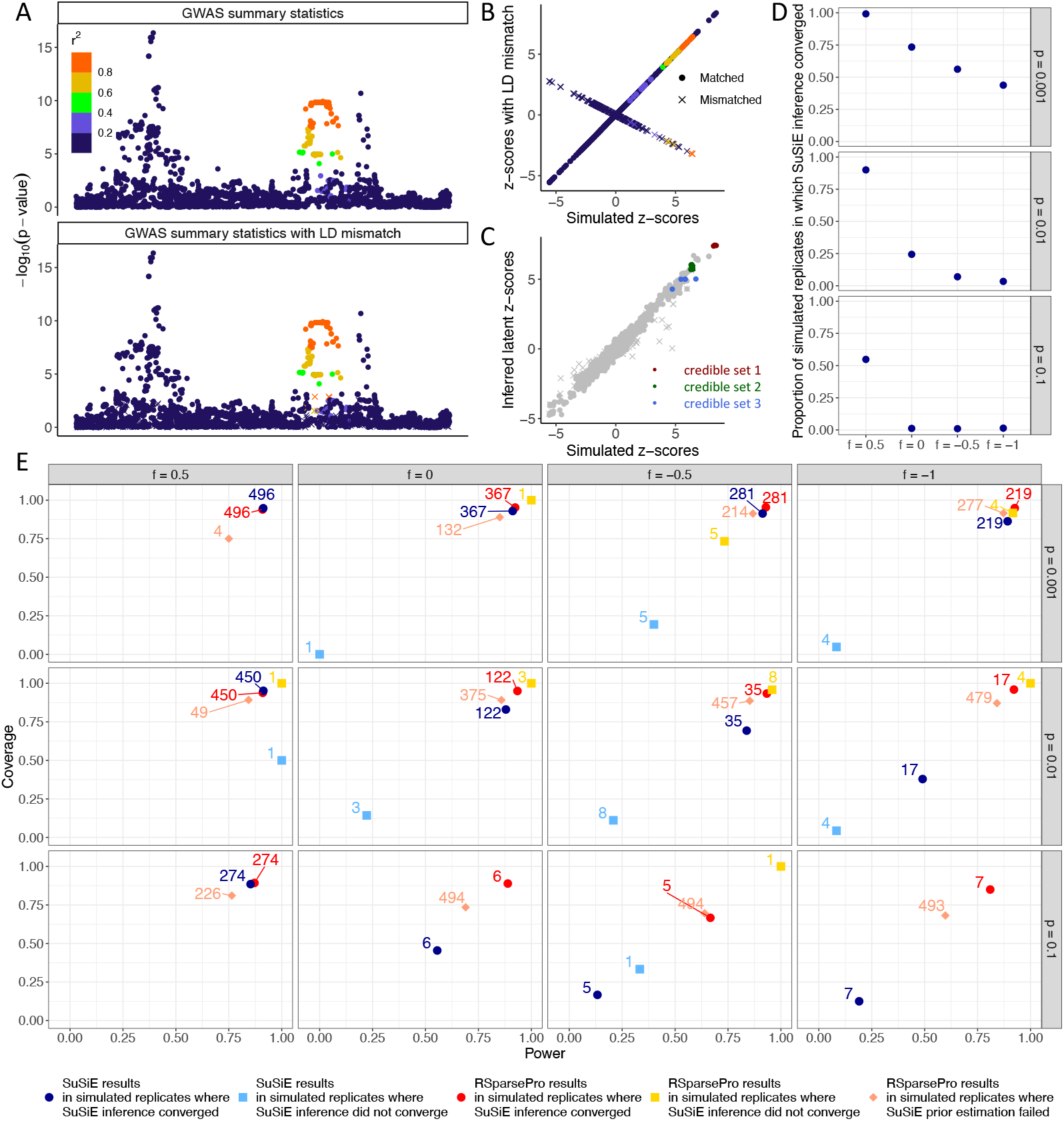
Robust fine-mapping in simulation studies with *K*_*C*_ = 3 causal variants. (A) Exemple locus with three causal variants and 10% of the variants having a mismatch factor of -0.5 (*K*_*c*_ = 3, *p* = 0.1, *f* = −0.5). The top panel displays the statistical significance of each variant based on GWAS summary statistics without LD mismatch. The bottom panel displays the statistical significance of each variant based on GWAS summary statistics with LD mismatch. Each dot represents a variant not subject to mismatch while each cross represents a variant affected by LD mismatch. The variants are colored by the magnitude of correlation (LD *r*^2^) with one of the simulated causal variants. (B) Comparison of simulated z-scores without LD mismatch and z-scores with LD mismatch in the example. (C) Comparison of simulated z-scores with LD mismatch and the latent z-scores inferred by RSparsePro. RSparsePro used the z-scores with LD mismatch as input in the example. Variants in credible sets identified by RSparsePro are indicated. (D) Proportion of simulated replicates where the SuSiE inference algorithm converged. Rows correspond to different proportions of mismatched variants. Columns correspond to different mismatch factors. (E) Power and coverage of RSparsePro and SuSiE across various simulation settings. The simulated replicates were categorized based on whether SuSiE encountered convergence issues. SuSiE does not provide fine-mapping results when its prior estimation fails. The numbers of simulated replicates in each category are labelled, totaling 500 in each simulation setting for each method.

Across all simulation settings, as expected, SuSiE frequently encountered convergence issues when the proportion of mismatched variants or the magnitude of LD mismatch was high (**Figure** 2D and **Supplementary Table** S1). In simulation replicates where its inference algorithm did not converge, SuSiE may provide redundant credible sets that did not contain causal variants (**Supplementary Figure** S1 and **Supplementary Table** S1) and could have substantially reduced power and coverage (**Figure** 2E).

RSparsePro consistently demonstrated comparable power and coverage as SuSiE when there was no or minimal LD mismatch (**Supplementary Table** S2). For example, when the z-scores of 0.1% of the variants were multiplied by a mismatch factor of 0.5 (*K*_*c*_ = 3, *p* = 0.001, *f* = 0.5), the SuSiE inference algorithm successfully converged in 496 of the 500 simulated replicates (**Figure** 2E and **Supplementary Table** S2). In these 496 simulated replicates, RSparsePro achieved a power of 0.909 and a coverage of 0.940, compared to SuSiE with a power of 0.913 and a coverage of 0.948 (**Figure** 2E and **Supplementary Table** S2).

With increased magnitude of LD mismatch (*K*_*c*_ = 3, *p* = 0.001, *f* = −1.0), the SuSiE inference algorithm converged in 219 simulated replicates (**Figure** 2E and **Supplementary Table** S2). Among these 219 simulated replicates, RSparsePro achieved a power of 0.925 and a coverage of 0.949 while SuSiE achieved a power of 0.892 and a coverage of 0.863 (**Figure** 2E and **Supplementary Table** S2).

With increased proportion of mismatched variants, prior estimation in SuSiE was more likely to fail. For instance, with 10% of the variants having a mismatch factor of −1.0 (*K*_*c*_ = 3, *p* = 0.1, *f* = −1.0), prior estimation in SuSiE failed in 493 of the 500 simulated replicates, while RSparsePro achieved a power of 0.598 and a coverage of 0.682 in the same simulated replicates (**Figure** 2Eand **Supplementary Table** S2).

As expected, these trends were consistent with increased number of simulated causal variants (**Supplementary Figures** S2-4).

### 2.3 Fine-mapping of cis-pQTL summary statistics

To evaluate the performance of RSparsePro in real data, we first performed fine-mapping using cis-protein quantitative trait loci (cis-pQTLs) summary statistics from the Fenland and the deCODE proteogenomic studies (**Methods**), both including participants of European ancestry [14, 15]. Since individual-level data were not available from these two studies, we used 5,000 randomly selected European ancestry individuals from the UK Biobank [21] as the LD reference panel. Notably, the Fenland study was based in England, where the population is more likely to share similar LD structures with the UK Biobank LD reference panel. In contrast, the deCODE study was based in Iceland, where genetic drift in recent demographic history may have resulted in more frequent LD discrepancies between the GWAS population and the reference panel [22].

As expected, for the 1,353 proteins for which at least one genome-wide significant cis-genetic association was identified in both studies, SuSiE was more likely to encounter convergence issues in the deCODE study (**Figure** 3A and **Supplementary Tables** S3-4). Specifically, prior estimation in SuSiE failed in 84 (6.2%) cis-pQTLs in the Fenland study and 259 (19.1%) in the deCODE study. Additionally, the SuSiE inference algorithm did not converge in 128 (9.5%) cis-pQTLs in the Fenland study and 187 (13.8%) cis-pQTLs in the deCODE study (**Figure** 3A and **Supplementary Tables** S3-4). Consistent with findings from simulation studies, in cis-pQTLs where the inference algorithm did not converge, SuSiE tended to identify a large number of credible sets, with a median of 9 (IQR: 8-10) in the Fenland study (vs. median = 2, IQR: 1-4 in cis-pQTLs when the algorithm converged) and 9 (IQR: 8-10) in the deCODE study (vs. median = 5, IQR: 2-9 in cis-pQTLs when the algorithm converged; **Figure** 3B and **Supplementary Tables** S3-4). In contrast, the distributions of the number of credible sets identified by RSparsePro remained largely consistent in cis-pQTLs where SuSiE encountered convergence issues (**Figure** 3B and **Supplementary Tables** S3-4).

**Figure 3.**
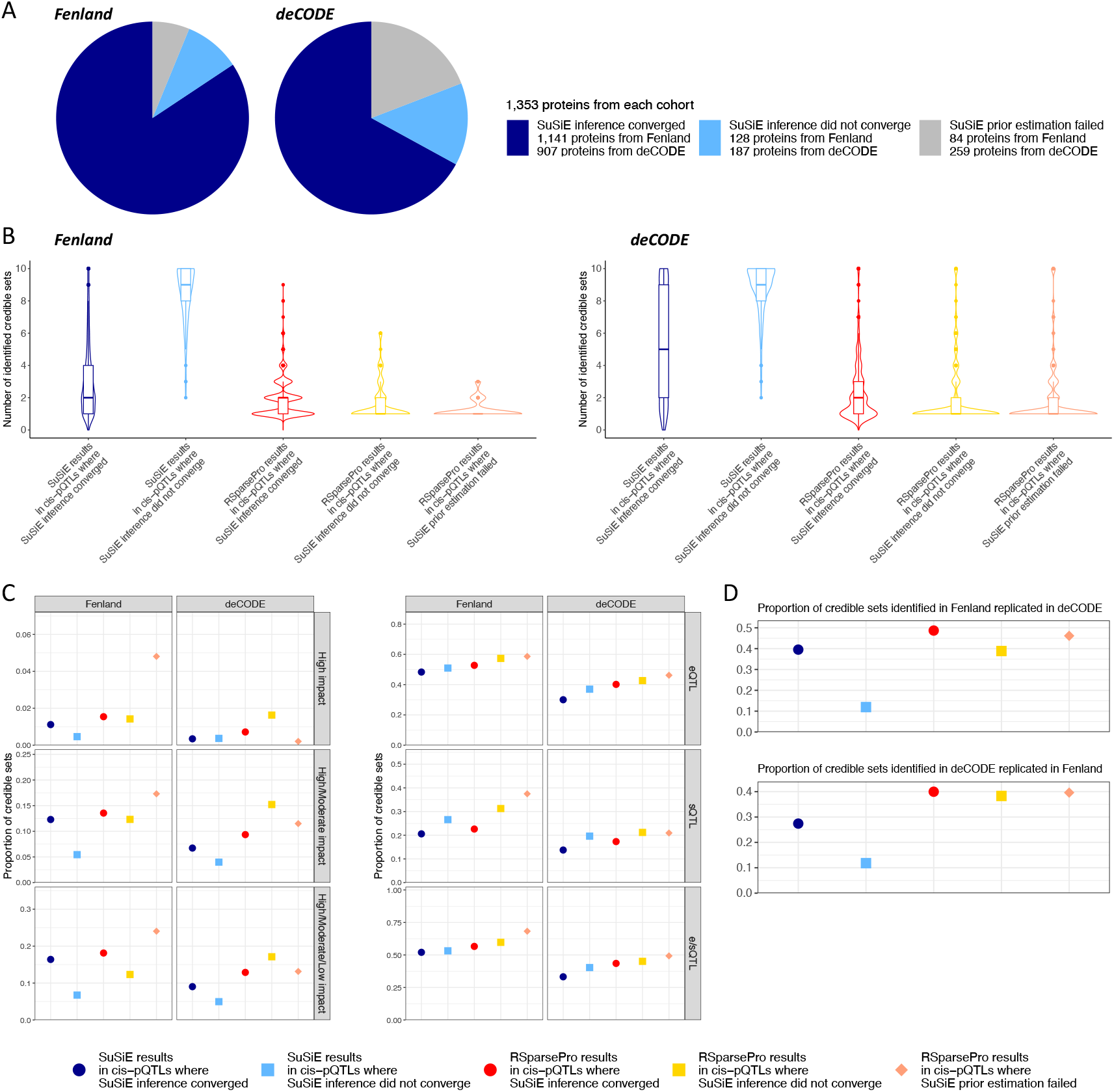
Summary of fine-mapping results for 1,353 plasma proteins with a cis-protein quantitative trait locus (cispQTL). (A) Proportions of proteins for which SuSiE encountered convergence issues. Numbers of cis-pQTLs where the SuSiE inference algorithm converged, where the SuSiE inference algorithm did not converge, and where prior estimation in SuSiE failed are indicated. (B) Distributions of numbers of credible sets identified by SuSiE and RSparsePro. The cis-pQTLs are categorized based on whether SuSiE encountered convergence issues. For each box plot, the edges represent the first and third quartiles, the line inside the box represents the median value, and the whiskers extend to the smallest value within 1.5 times the inter-quartile range below the first quartile and the largest value within 1.5 times the inter-quartile range above the third quartile. Dots represent outliers. The shape of each violin plot indicates the density of data. (C) Proportions of credible sets containing variants with predicted functional impacts or potential regulatory roles in gene expression. The proportions of credible sets containing at least one variant with a (1) high impact, (2) high or moderate impact, or (3) high, moderate, or low impact on the corresponding protein-coding genes are indicated. The proportions of credible sets containing at least one variant being (4) an expression quantitative trait locus (eQTL), (5) a splicing quantitative trait locus (sQTL), or (6) eQTL or sQTL (e/sQTL) of the corresponding protein-coding genes in at least one GTEx tissue are indicated. (D) Replication rates of credible sets. The proportions of credible sets identified in one study that shared at least one variant with credible sets identified in the other study are indicated.

In both the Fenland and deCODE studies, RSparsePro robustly identified credible sets that were more likely to include variants with functional impacts on protein-coding genes or potential regulatory roles in gene expression (**Figure** 3C and **Supplementary Tables** S5-8). For example, in cis-pQTLs where the SuSiE inference algorithm converged in the Fenland study, 290 (13.6%) of the 2,140 RSparsePro-identified credible sets contained at least one variant with an Ensembl Variant Effect Predictor (VEP [23, 24])-predicted high or moderate functional impact on the corresponding protein-coding genes, similar to 12.3% using SuSiE (Chi-squared test p-value = 0.18; **Figure** 3C). In cis-pQTLs where the SuSiE inference algorithm did not converge, 26 (12.3%) of the 211 RSparsePro-identified credible sets contained at least one variant with a VEP-predicted high or moderate functional impact, significantly outperforming SuSiE (5.4%, Chi-squared test p-value = 3.7 × 10^−4^; **Figure** 3C). In cis-pQTLs where prior estimation in SuSiE failed, 18 (17.3%) of the 104 credible sets identified by RSparsePro contained at least one variant with a VEP-predicted high or moderate functional impact (**Figure** 3C), whereas SuSiE was unable to provide results. Furthermore, based on the Genotype-Tissue Expression (GTEx) dataset [25], in both studies, RSparsePro consistently identified more credible sets with at least one variant significantly associated with expression levels or alternative splicing of the corresponding protein-coding genes compared to SuSiE (**Figure** 3C and **Supplementary Tables** S3-4).

Importantly, credible sets identified by RSparsePro demonstrated a higher cross-study replication rate than those identified by SuSiE (**Figure** 3D and **Supplementary Tables** S3-4). Specifically, in the Fenland study, in cis-pQTLs where the SuSiE inference algorithm converged, 1,042 (48.7%) of the 2,140 RSparsePro-identified credible sets were replicated in the deCODE study (vs. 39.5% using SuSiE; Chi-squared test p-value = 1.6 × 10^−11^; **Figure** 3D). In loci where the SuSiE inference algorithm did not converge or the prior estimation in SuSiE failed, RSparsePro achieved replication rates of 38.9% and 46.2%, respectively (**Figure** 3D). In comparison, SuSiE had a replication rate of 11.9% (Chi-squared test p-value = 9.0 × 10^−22^) in cis-pQTLs where the inference algorithm did not converge. On the other hand, credible sets identified in the deCODE study using both methods had a lower replication rate in the Fenland study (**Figure** 3D). Nonetheless, RSparsePro achieved replication rates of 40.0%, 38.3%, and 39.6% in cis-pQTLs where the SuSiE inference algorithm converged (vs. 27.4% using SuSiE; Chi-squared test p-value = 3.3 × 10^−25^), where the SuSiE inference algorithm did not converge (vs. 11.8% using SuSiE; Chi-squared test p-value = 5.0 × 10^−44^), and where prior estimation in SuSiE failed, respectively (**Figure** 3D).

RSparsePro enabled robust fine-mapping of cis-pQTLs for proteins with important biological functions. For example, PCSK9 plays a crucial role in cholesterol metabolism by regulating the clearance of circulating LDL cholesterol through its binding to LDL receptors and has been a major target for cholesterol-lowering drugs [26, 27, 28, 29, 30]. In the Fenland study, both SuSiE (**Figure** 4A) and RSparsePro (**Figure** 4B) identified the same three credible sets based on cis-pQTL summary statistics for plasma PCSK9. In the deCODE study, likely due to LD mismatch, prior estimation in SuSiE failed. In contrast, RSparsePro replicated all three credible sets identified in the Fenland study and additionally identified one credible set in which the variants were suggestively but not genome-wide significant in the Fenland study (**Figure** 4C), highlighting the robustness of RSparsePro in the presence of complex LD structures.

**Figure 4.**
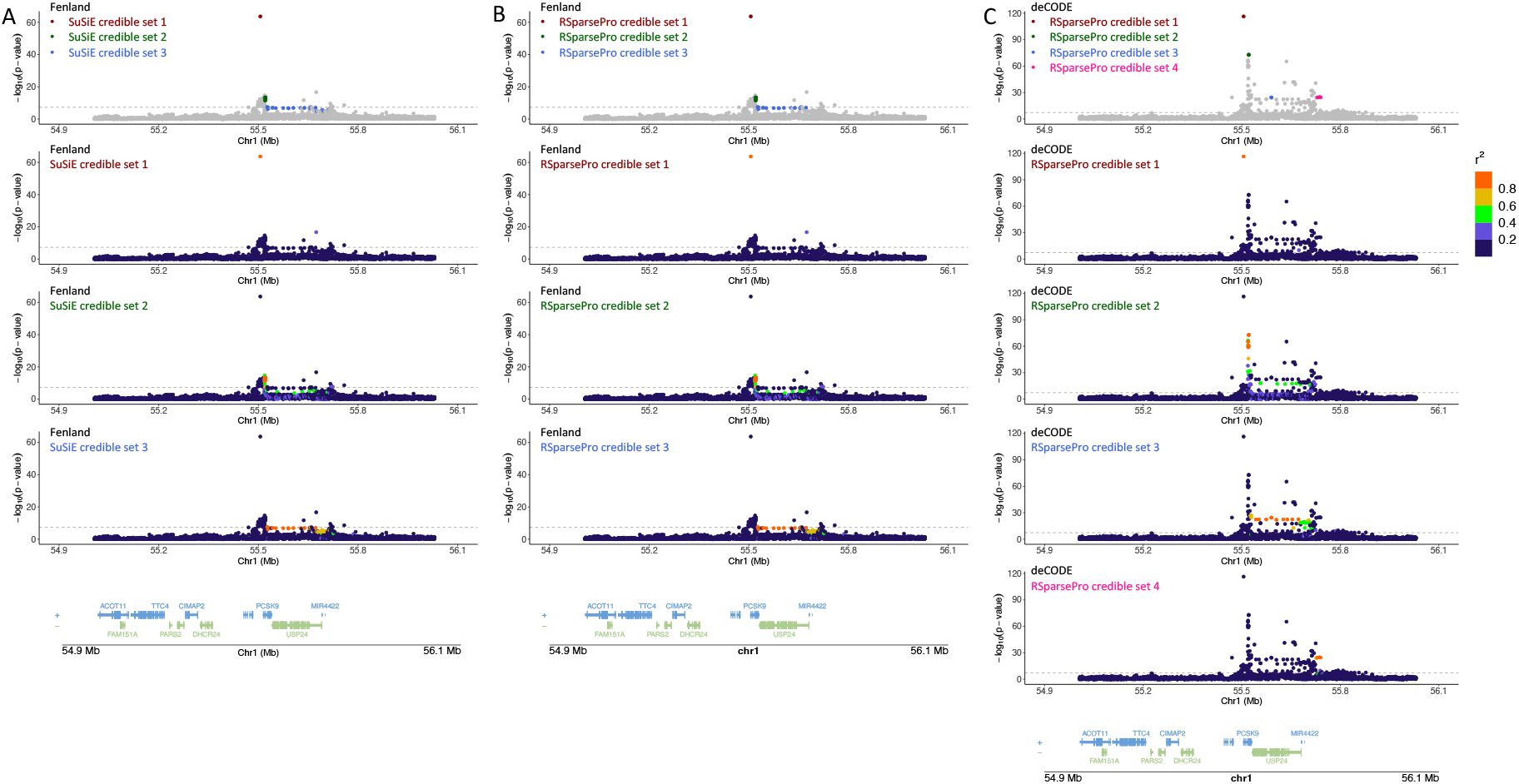
Fine-mapping of cis-protein quantitative trait locus for plasma PCSK9. Fine-mapping results using (A) SuSiE in the Fenland study, (B) RSparsePro in the Fenland study, and (C) RSparsePro in the deCODE study are illustrated. The identified credible sets are indicated. Genetic variants in this locus are plotted with their significance in respective studies, and colored by the magnitude of correlation (LD *r*^2^) with the lead variant in each credible set. The UCSC known gene tracks are presented, with gene models colored by their respective strands. SuSiE could not provide fine-mapping results in the deCODE study due to failed prior estimation.

### 2.4 Fine-mapping of ancestry-specific LDL cholesterol GWAS summary statistics

Next, we evaluated the performance of RSparsePro in fine-mapping large-scale meta-analyses of GWAS for which in-sample LD is unavailable. We utilized ancestry-specific LDL cholesterol GWAS summary statistics from the Global Lipids Genetics Consortium (GLGC) [31] and individuals with similar genetic ancestries from the 1000 Genomes Project [32] as LD reference panels (**Methods**). SuSiE encountered convergence issues in 60 (12.2%) of the 491 genome-wide significant loci across the five ancestry-specific GWAS, including 55 (11.2%) loci where prior estimation failed and 5 (1.0%) loci where the SuSiE inference algorithm did not converge (**Figure** 5A). Similar to the cis-pQTL analyses, in loci where the SuSiE inference algorithm did not converge, SuSiE identified more credible sets (median = 8; IQR: 5-9) than in loci where the algorithm converged (median = 1; IQR: 1-2; **Figure** 5B and **Supplementary Table** S9). In contrast, the distributions of the number of credible sets identified by RSparsePro remained largely consistent (**Figure** 5B and **Supplementary Table** S9).

**Figure 5.**
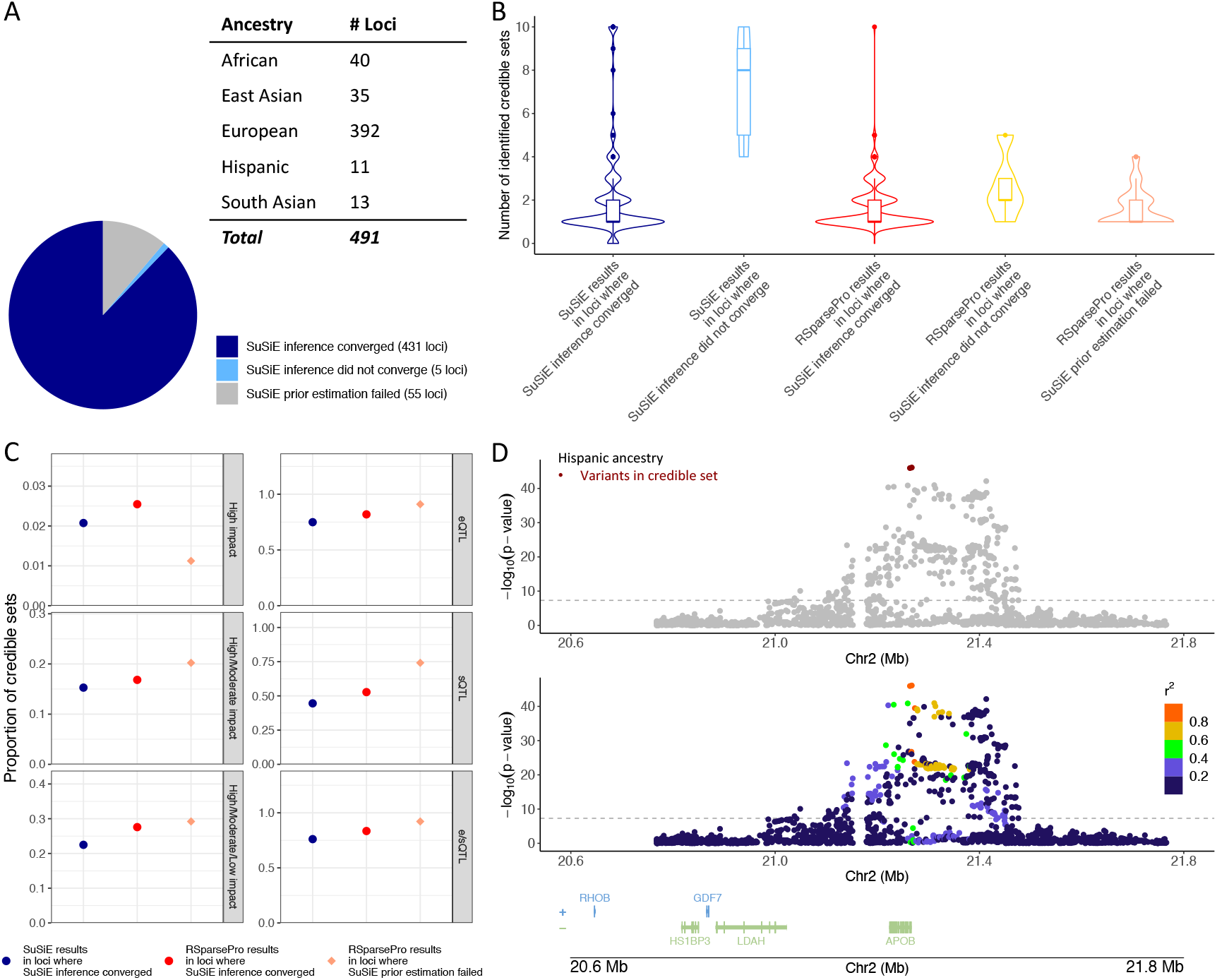
Summary of fine-mapping results for genome-wide significant loci in ancestry-specific GWAS of LDL cholesterol. (A) Proportions of loci for which SuSiE encountered convergence issues. Numbers of genome-wide significant loci identified in each ancestry-specific meta-analysis are indicated. Numbers of loci where the SuSiE inference algorithm converged, where the SuSiE inference algorithm did not converge, and where prior estimation in SuSiE failed are indicated. (B) Distributions of numbers of credible sets identified by SuSiE and RSparsePro. The loci are categorized based on whether SuSiE encountered convergence issues. For each box plot, the edges represent the first and third quartiles, the line inside the box represents the median value, and the whiskers extend to the smallest value within 1.5 times the inter-quartile range below the first quartile and the largest value within 1.5 times the inter-quartile range above the third quartile. Dots represent outliers. The shape of each violin plot indicates the density of data. (C) Proportions of credible sets containing variants with predicted functional impacts or potential regulatory roles in gene expression. The proportions of credible sets containing at least one variant with a (1) high impact, (2) high or moderate impact, or (3) high, moderate, or low impact on any genes are indicated. The proportions of credible sets containing at least one variant being (4) an expression quantitative trait locus (eQTL), (5) a splicing quantitative trait locus (sQTL), or (6) eQTL or sQTL (e/sQTL) of any genes in at least one GTEx tissue are indicated. Results for loci where the SuSiE inference algorithm did not converge are not displayed because of the small number of loci in this category, and are available in **Supplementary Table** S9. (D) Fine-mapping of the *APOB* locus based on the Hispanic ancestry-specific GWAS. The RSparsePro-identified credible set is indicated. Genetic variants in this locus are plotted with their significance in the GWAS, and colored by the magnitude of correlation (LD *r*^2^) with the lead variant in the credible set. The UCSC known gene tracks are presented, with gene models colored by their respective strands.

Credible sets identified by RSparsePro contained more variants with functional impacts or potential regulatory roles in gene expression (**Figure** 5C and **Supplementary Tables** S10-11). For example, across all ancestries, in loci where the SuSiE inference algorithm converged, RSparsePro identified a total of 707 credible sets. Of these, 119 (16.8%) contained at least one variant with a VEP-predicted high or moderate functional impact on any gene, while 590 (83.5%) contained at least one variant significantly associated with expression levels or alternative splicing of at least one gene in at least one GTEx tissue, compared to 15.3% (Chi-squared test p-value = 0.44) and 76.1% (Chi-squared test p-value = 3.6 × 10^−4^) using SuSiE, respectively (**Figure** 5C). In loci where prior estimation in SuSiE failed, 18 (20.2%) of the 89 credible sets identified by RSparsePro contained at least one variant with a VEP-predicted high or moderate functional impact, and 82 (92.1%) contained at least one variant significantly associated with expression levels or alternative splicing (**Figure** 5C). The robust performance of RSparsePro was consistent across the five ancestries, although the majority of loci were identified in the European ancestry-specific GWAS (**Supplementary Table** S9).

RSparsePro identified potential causal variants in genes crucial for lipid metabolism. For instance, based on the Hispanic ancestry-specific GWAS, one credible set was identified in *APOB* (**Figure** 5D), which encodes the main protein component of LDL. This credible set contained a missense variant, rs1367117 (p.Thr98Ile), which has been associated with various lipid biomarkers in large-scale GWAS across multiple ancestries [31, 33, 34] and demonstrated a potential maternal-specific association with adiposity in family-based studies [35]. In contrast, prior estimation in SuSiE failed for this locus, likely due to LD discrepancies between the Hispanic ancestry GWAS populations and the Admixed American ancestry population from the 1000 Genomes Project used as the LD reference panel.

Moreover, RSparsePro facilitated the interpretation of causal variants and their regulatory mechanisms (**Supplementary Table** S12). For instance, one credible set was identified in *GCKR* based on the European ancestry-specific GWAS (**Figure** 6A). This credible set contained a missense variant, rs1260326 (p.Leu446Pro), located in a ChromHMM-predicted enhancer in the liver [36, 37] and in a Capture-C-detected promoter-interacting region in the HepG2 cell line mapping to *GCKR* [38]. These findings support the causal effect of the variant, possibly indirectly through regulating glucokinase activity and carbohydrate metabolism, primarily in the liver [39, 40] (**Supplementary Table** S12). Two credible sets were identified in the *RHCE-MACO1-LDLRAP1* locus based on the same GWAS (**Figure** 6B). Interestingly, while both credible sets contained variants in promoter-interacting regions mapping to *RHCE*, one may simultaneously regulate the expression of *LDLRAP1* (**Supplementary Table** S12). Although *RHCE* is known to have strong pleiotropic effects [41, 42, 43], its association with lipid levels has not been fully elucidated. In contrast, *LDLRAP1* encodes an adaptor protein involved in the endocytosis of LDL particles through interaction with LDL receptors [44, 45], which directly supports the regulatory role of the identified variants in lipid metabolism. Furthermore, one credible set was identified in the *SLC17A8-NR1H4* locus based on the East Asian ancestry-specific GWAS, where multiple variants in the credible set were located in ChromHMM-predicted enhancers in the liver and promoter-interacting regions in the HepG2 cell line mapping to *NR1H4* (**Figure** 6C and **Supplementary Table** S12). Given that *NR1H4* is a bile acid receptor and master regulator of genes involved in bile acid synthesis and transport [46, 47, 48], which are closely linked to lipid homeostasis, the functional relevance of these variants further strengthens the evidence for their involvement in lipid metabolism. Notably, prior estimation in SuSiE failed in all of these loci, likely due to LD mismatch. These results highlight the broad utility of RSparsePro in characterizing biologically relevant associations and improving variant-to-gene mapping.

**Figure 6.**
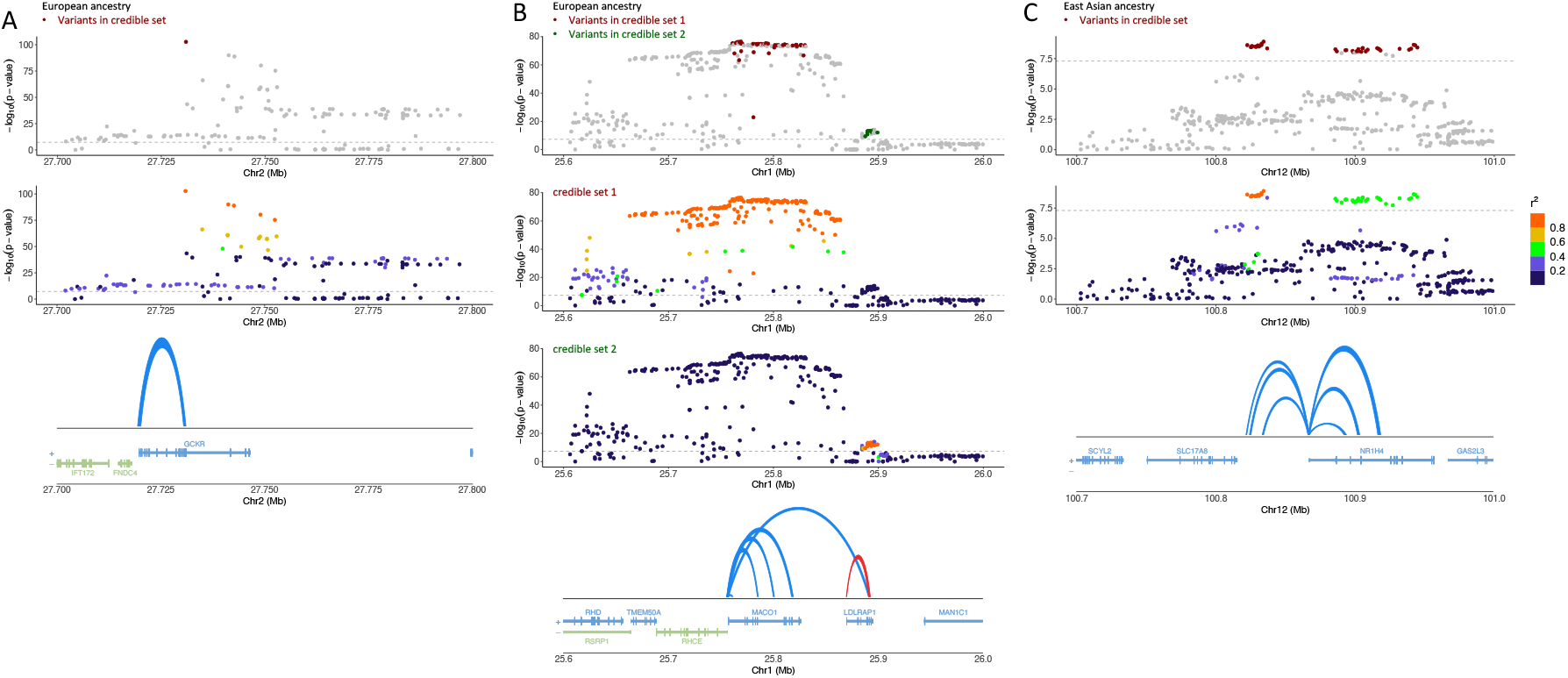
Exemple loci where RSparsePro identified potential regulatory elements for LDL cholesterol. Fine-mapping results obtained using RSparsePro for (A) the *GCKR* locus based on the European ancestry-specific GWAS, (B) the *RHCE-MACO1-LDLRAP1* locus based on the European ancestry-specific GWAS, and (C) the *SLC17A8-NR1H4* locus based on the East Asian ancestry-specific GWAS are illustrated. The identified credible sets are indicated. Genetic variants in each locus are plotted with their significance in respective ancestry-specific GWAS, and colored by the magnitude of correlation (LD *r*^2^) with the lead variant in each credible set. The UCSC known gene tracks are presented, with gene models colored by their respective strands. Promoter regions and promoter-interacting regions detected in Capture-C data in the HepG2 cell line are connected by arches. Coordinates of promoter-interacting regions overlapping with the identified credible sets and the corresponding promoter regions are available in **Supplementary Table** S12. SuSiE could not provide fine-mapping results in these loci due to failed prior estimation.

## 3 Discussion

Accurate identification of potential causal variants through fine-mapping is crucial for understanding the genetic architecture of human traits and diseases, as well as for informing functional follow-up studies [3, 6]. One of the most pressing challenges hindering the broad application of fine-mapping is the unavailability of in-sample LD information and the inconsistencies between GWAS populations and LD reference panels. In this work, we present RSparsePro, a method designed for robust fine-mapping in the presence of potential LD mismatch, directly addressing this challenge.

Instead of directly modeling the z-scores from GWAS summary statistics, we introduced a latent variable to help mitigate the impact of LD mismatch. This unique feature is the key to the robustness of RSparsePro and serves as the primary factor contributing to its improved performance compared to SuSiE. This approach may also be adapted for other tasks in statistical genetics research that rely on GWAS summary statistics and LD reference panels. Additionally, the use of an uninformative prior for the causal effect sizes eliminates LD mismatch-induced issues in prior estimation.

Through extensive simulation studies, we demonstrated that RSparsePro achieved comparable power and coverage as SuSiE in identifying true causal variants when there was no or minimal LD mismatch. However, as the proportion and magnitude of LD mismatch increased, SuSiE had a dramatically higher frequency of convergence issues and could identify redundant credible sets that do not contain true causal variants when its inference algorithm did not converge. Notably, even when the SuSiE inference algorithm did converge, the credible sets identified could still have compromised power and coverage. In contrast, RSparsePro maintained largely consistent performance across loci where the SuSiE inference algorithm converged, those where the SuSiE inference algorithm did not converge, and those where prior estimation in SuSiE failed.

These trends were also observed in real data analyses involving fine-mapping based on cis-pQTL summary statistics and large-scale GWAS meta-analysis summary statistics, respectively. Although in-sample LD information was unavailable for these studies, RSparsePro consistently identified credible sets with lower redundancy and a stronger enrichment of variants having predicted functional impacts or potential regulatory roles in gene expression. Additionally, despite potential population-specific genetic effects, the consistently higher cross-study replication rates of credible sets identified in cis-pQTLs using RSparsePro supported the validity of our findings.

We demonstrated that fine-mapping with RSparsePro can confirm potential causal variants implicating biologically relevant genes, such as *APOB*, a clinically actionable gene for familial hyperlipidemia [49, 50]. This locus, identified in a Hispanic ancestry-specific GWAS for LDL cholesterol, could not be fine-mapped using SuSiE with the 1000 Genomes Project Admixed American ancestry reference population. Furthermore, combined with functional evidence, fine-mapping using RSparsePro may further suggest potential regulatory mechanisms and improve variant-to-gene mapping, such as through the identification of credible sets overlapping with enhancers and promoter-interacting regions. These results underscore the utility of RSparsePro in facilitating the biological interpretation of GWAS findings, especially in loci containing multiple genes with complex LD structures.

It is important to note that while RSparsePro offers robust fine-mapping results, its power and coverage can still decline substantially under widespread LD mismatch in simulations, although to a lesser extent than SuSiE. Based on cis-pQTL summary statistics, credible sets identified by RSparsePro in the deCODE study showed weaker enrichment in variants with predicted functional impacts and lower cross-study replication rates compared to those identified in the Fenland study. This discrepancy may arise from more frequent and severe LD mismatch between the Icelandic study population and the UK Biobank LD reference panel [22], although differences in sample size may also play a role. Therefore, in practice, it remains preferable to utilize an LD reference panel closely matching the GWAS population to enhance the reliability of fine-mapping results.

In summary, RSparsePro is a robust method for fine-mapping in the presence of potential LD mismatch. Its ability to identify causal variants across a varying degree of LD mismatch significantly enhances the applicability of fine-mapping analyses, particularly in the context of increasingly large GWAS that include protected datasets or involve multiple cohorts and diverse populations. Ultimately, RSparsePro has the potential to help advance the goal of translating genetic associations into biological insights and clinical applications.

## 4 Methods

### 4.1 Simulation study design

We conducted simulation studies with a varying degree of LD mismatch to evaluate the performance of RSparsePro and SuSiE. We obtained genotype information from 503 European ancestry individuals from the 1000 Genomes Project in 10 randomly sampled 1-megabase loci [32] and calculated the corresponding LD matrices (**Supplementary Table** S1).

In each locus, we randomly sampled *K*_*C*_ ∈ {3, 5} independent causal variants (LD |*r*| *<* 0.1) and followed previous studies [51, 52, 53] to simulate GWAS summary statistics z-scores with a targeted sample size (*N*_*target*_) of 500,000 and a per-variant heritability 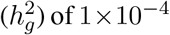 of 1×10^−4^. Specifically, we first simulated the observed marginal variant-phenotype associations

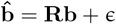

where **R** represents the LD matrix; **b** is a *G* × 1 vector representing the true causal effect sizes; and ϵ is a *G* × 1 vector representing the estimation errors. The estimation errors were simulated by first sampling a null phenotype for the 1000 Genomes Project European ancestry population, 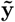, from 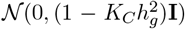. We regressed this null phenotype on the genotypes to obtain the marginal variant-null phenotype associations, 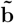, in the LD reference panel (*N*_0_ = 503). We then re-scaled the estimation errors to the targeted sample size

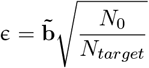

The simulated z-scores were obtained as

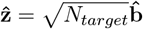

Next, to simulate a varying degree of LD mismatch, we randomly selected a proportion of non-causal variants, *p* ∈ {0.001, 0.01, 0.1} and multiplied their GWAS summary statistics z-scores by a mismatch factor of *f* ∈ {0.5, 0.0, −0.5, −1.0}. For example, with *p* = 0.1 and *f* = −1.0, 10% of the variants in a locus had their GWAS z-scores with a flipped sign. For each of the 10 loci, we generated 50 simulation replicates, totaling 500 replicates for each combination of the number of causal variants (*K*_*C*_), mismatch proportion (*p*) and mismatch factor (*f*).

For each simulated replicate, we performed fine-mapping using SuSiE and RSparsePro, respectively, with the LD matrix and the simulated z-scores as input. For SuSiE, we utilized the “susie rss” function with default settings. In both SuSiE and RSparsePro, the maximum number of credible sets was set to 10. We categorized the simulation replicates based on whether SuSiE encountered convergence issues: “SuSiE prior estimation failed”, “SuSiE inference did not converge” and “SuSiE inference converged”. Notably, when prior estimation in SuSiE fails, SuSiE does not provide fine-mapping results.

We compared the performance of SuSiE and RSparsePro based on three metrics:

1. Redundancy: We summarized the number of identified credible sets.
2. Power: We calculated the proportion of true causal variants included in the credible sets.
3. Coverage: We calculated the proportion of credible sets containing true causal variants.

### 4.2 Fine-mapping of cis-pQTL summary statistics

We obtained GWAS summary statistics for plasma protein levels from both the Fenland and deCODE studies [14, 15]. Details of these studies have been previously described [14, 15]. The Fenland study included 10,708 individuals of European ancestry residing in Cambridgeshire, England, at the time of recruitment [54], while the deCODE study included 35,559 individuals of European ancestry residing in Iceland at the time of recruitment [15]. Both studies measured plasma protein levels using the SomaLogic SomaScan v4 assay. To enable comparison of fine-mapping results between these two cohorts, we restricted our analyses to 1,353 plasma proteins for which at least one genome-wide significant cis-genetic association was identified [14, 15], located outside the major histocompatibility complex (MHC) region (GRCh37 chr6:28,477,797-33,448,354). These cis-variants were located within 500 kilobases of the gene body of the respective protein-coding genes.

For each protein, we performed fine-mapping based on GWAS summary statistics for cis-pQTLs, including all variants located within 500 kilobases of the gene body of the protein-coding gene, using SuSiE and RSparsePro separately with default settings and up to 10 credible sets. We utilized 5,000 randomly selected unrelated individuals from the UK Biobank as the LD reference panel [21]. We retained variants that were present in both GWAS populations, and that had a minor allele frequency (MAF) *>* 0.01 in the LD reference panel.

Similarly, we categorized the cis-pQTLs into three groups based on whether the SuSiE algorithm encountered convergence issues: “SuSiE prior estimation failed”, “SuSiE inference did not converge” and “SuSiE inference converged”. Within each category, we investigated three metrics for SuSiE and RSparsePro:

1. Redundancy: We summarized the number of identified credible sets.
2. Impact: We annotated the functional impact of each variant in the credible sets using the Ensembl VEP (release 112) [23, 24]. We calculated the proportion of credible sets containing at least one variant with a VEP-annotated high, moderate, or low functional impact on the corresponding protein-coding genes, as opposed to modifiers. Specifically, high-impact variants are predicted to have a disruptive effect on gene or protein structure and function; moderate-impact variants are missense mutations that do not necessarily lead to complete loss of function; low-impact variants may have limited, regulatory effects; and modifier variants are assumed to have minimal effects on gene function [23, 24]. We also calculated the proportion of credible sets containing at least one variant significantly associated with expression levels or alternative splicing, or both, of the corresponding protein-coding genes, with a tissue-specific false discovery rate (FDR) *<* 0.05 in any tissue available in the GTEx dataset (version 8) [25].
3. Replication rate: We calculated the proportion of credible sets identified in one study that shared at least one variant with credible sets identified in the other study for the same protein.

### 4.3 Fine-mapping of ancestry-specific LDL GWAS summary statistics

We obtained ancestry-specific GWAS summary statistics for LDL cholesterol from the GWAS meta-analysis conducted by the GLGC [31]. Details of the participating studies and the meta-analysis have been previously described [31]. The meta-analysis included up to 99,432 individuals of Admixed African or African (referred to as “African”) ancestry, 146,492 individuals of East Asian ancestry, 1,320,016 individuals of European ancestry, 48,057 individuals of Hispanic ancestry, and 40,963 individuals of South Asian ancestry. For each ancestry-specific GWAS, we defined a genome-wide significant locus as a ±500-kilobase region around the variant showing the strongest statistical significance (p-value *<* 5 × 10^−8^) within this region. We excluded loci that overlapped with the MHC region.

For each ancestry-specific GWAS, we performed fine-mapping of each locus using SuSiE and RSparsePro separately, with default settings and up to 10 credible sets. We used the African (N = 661), East Asian (N = 504), European (N = 503), Admixed American (N = 347, as LD reference panel for the Hispanic ancestry-specific GWAS), and South Asian (N = 489) ancestry populations from the 1000 Genomes Project [32] as LD reference panels for the respective ancestry-specific GWAS. Given the small sample sizes of the LD reference panels, we retained only variants with a MAF *>* 0.05 in the corresponding LD reference panel for each ancestry-specific GWAS.

Again, we categorized the loci into three groups based on whether the SuSiE algorithm encountered convergence issues: “SuSiE prior estimation failed”, “SuSiE inference did not converge” and “SuSiE inference converged”. Within each category, we investigated the metrics of redundancy and impact for SuSiE and RSparsePro, as described above, except that variants in credible sets may have functional impacts on any genes or be associated with expression levels or alternative splicing of any genes.

Furthermore, we conducted functional genomic annotation for all variants in the identified credible sets. Specifically, we utilized the pre-trained ChromHMM core 15-state model to predict the chromatin state of each variant in the liver (E066) and adipose tissue (E063) [36, 37]. ChromHMM uses chromatin marks, primarily histone modifications and chromatin accessibility in biopsy samples or cell lines, to identify recurrent patterns (i.e., chromatin states) across the genome, including enhancers [36, 37]. Additionally, we used promoter-focused Capture-C data from the HepG2 hepatoblastoma cell line and the *in vitro*-differentiated SGBS preadipocyte cell line to identify promoter-interacting regions that are potential regulatory elements [38]. Significant promoter-interacting regions were defined by a CHiCAGO score ≥ 5 at restriction fragment resolution [38]. For each variant in the identified credible sets, we assessed whether it was located in a promoter-interacting region in either the HepG2 cell line or the differentiated SGBS cell line and annotated the corresponding gene promoter. Variants located in credible sets that overlapped with ChromHMM-predicted enhancers and Capture-C-detected promoter-interacting regions were considered to have strong evidence supporting their role in regulating the expression of the corresponding genes.

## 5 Data Availability

The pQTL summary statistics from the Fenland study and the deCODE study were obtained from https://www.synapse.org/Synapse:syn51761394/wiki/622766 and https://www.decode.com/summarydata/, respectively. The ancestry-specific LDL cholesterol GWAS summary statistics were obtained from the GLGC https://csg.sph.umich.edu/willer/public/glgc-lipids2021/results/ancestry_specific/. VEP is available at https://useast.ensembl.org/info/docs/tools/vep/index.html. GTEx datasets were obtained from https://www.gtexportal.org/home/downloads/adult-gtex/qtl. ChromHMM chromatin state annotations were obtained from https://egg2.wustl.edu/roadmap/web_portal/chr_state_learning.html#core_15state. Promoter-focused Capture-C data were obtained from the Gene Expression Omnibus (GEO) with accession number GSE262496. The 1000 Genomes Project data are publicly available https://www.internationalgenome.org/. The UK Biobank data are available upon successful project application to the research committee https://www.ukbiobank.ac.uk/.

## 6 Software Availability

The RSparsePro software for robust fine-mapping is openly available at https://github.com/zhwm/RSparsePro_LD. The analysis conducted in this study is available at https://github.com/zhwm/RSparsePro_LD_analysis. SuSiE (version 0.12.35) was installed from CRAN at https://cran.r-project.org/web/packages/susieR/index.html.

## 7 Acknowledgements

W.Z. is supported by an Institut de valorisation des données (IVADO) Postdoctoral Fellowship, which is funded by the Canada First Research Excellence Fund. T.L. is supported by start-up funding from the Office of the Vice Chancellor for Research and Graduate Education, School of Medicine and Public Health, and Department of Population Health Sciences at the University of Wisconsin-Madison. G.L. is funded by the Canadian Institutes of Health Research (Projects 426541 and 486808), the Canada Research Chair Program, the Montreal Heart Institute Foundation, and the NIH/NHGRI (5UM1HG0120-04). This research has been conducted using the UK Biobank Resource under Application Number 62518.

## References

[1] Benner, C. et al. Finemap: efficient variable selection using summary data from genome-wide association studies. Bioinformatics 32, 1493–1501 (2016).

[2] Zou, Y., Carbonetto, P., Wang, G. & Stephens, M. Fine-mapping from summary data with the “sum of single effects” model. PLOS Genetics 18, e1010299 (2022).

[3] Zhang, W., Najafabadi, H. & Li, Y. Sparsepro: An efficient fine-mapping method integrating summary statistics and functional annotations. PLOS Genetics 19, e1011104 (2023).

[4] Hormozdiari, F., Kostem, E., Kang, E. Y., Pasaniuc, B. & Eskin, E. Identifying causal variants at loci with multiple signals of association. Genetics 198, 497–508 (2014).

[5] Chen, W. et al. Fine mapping causal variants with an approximate bayesian method using marginal test statistics. Genetics 200, 719–736 (2015).

[6] Wang, G., Sarkar, A., Carbonetto, P. & Stephens, M. A simple new approach to variable selection in regression, with application to genetic fine mapping. Journal of the Royal Statistical Society Series B: Statistical Methodology 82, 1273–1300 (2020).

[7] Kichaev, G. et al. Integrating functional data to prioritize causal variants in statistical fine-mapping studies. PLOS Genetics 10, e1004722 (2014).

[8] Hutchinson, A., Watson, H. & Wallace, C. Improving the coverage of credible sets in bayesian genetic fine-mapping. PLOS Computational Biology 16, e1007829 (2020).

[9] Wen, X., Lee, Y., Luca, F. & Pique-Regi, R. Efficient integrative multi-snp association analysis via deterministic approximation of posteriors. The American Journal of Human Genetics 98, 1114–1129 (2016).

[10] Newcombe, P. J., Conti, D. V. & Richardson, S. Jam: a scalable bayesian framework for joint analysis of marginal snp effects. Genetic Epidemiology 40, 188–201 (2016).

[11] Sesia, M., Katsevich, E., Bates, S., Candès, E. & Sabatti, C. Multi-resolution localization of causal variants across the genome. Nature Communications 11, 1093 (2020).

[12] Chen, W. et al. Improved analyses of gwas summary statistics by reducing data heterogeneity and errors. Nature Communications 12, 7117 (2021).

[13] Kanai, M. et al. Meta-analysis fine-mapping is often miscalibrated at single-variant resolution. Cell Genomics 2 (2022).

[14] Pietzner, M. et al. Mapping the proteo-genomic convergence of human diseases. Science 374, eabj1541 (2021).

[15] Ferkingstad, E. et al. Large-scale integration of the plasma proteome with genetics and disease. Nature Genetics 53, 1712–1721 (2021).

[16] Aragam, K. G. et al. Discovery and systematic characterization of risk variants and genes for coronary artery disease in over a million participants. Nature Genetics 54, 1803–1815 (2022).

[17] Zhang, W. et al. Sharepro: an accurate and efficient genetic colocalization method accounting for multiple causal signals. Bioinformatics 40, btae295 (2024).

[18] Zhang, W., Sladek, R., Li, Y., Najafabadi, H. S. & Dupuis, J. Accounting for genetic effect heterogeneity in finemapping and improving power to detect gene-environment interactions with sharepro. bioRxiv 2023–07 (2023).

[19] Blei, D. M., Kucukelbir, A. & McAuliffe, J. D. Variational inference: A review for statisticians. Journal of the American Statistical Association 112, 859–877 (2017).

[20] Titsias, M. & Lazaro-Gredilla, M. Spike and slab variational inference for multi-task and multiple kernel learning. Advances in Neural Information Processing Systems 24, 2339–2347 (2011).

[21] Bycroft, C. et al. The uk biobank resource with deep phenotyping and genomic data. Nature 562, 203–209 (2018).

[22] Ebenesersdóttir, S. S. et al. Ancient genomes from iceland reveal the making of a human population. Science 360, 1028–1032 (2018).

[23] McLaren, W. et al. The ensembl variant effect predictor. Genome Biology 17, 1–14 (2016).

[24] Cunningham, F. et al. Ensembl 2022. Nucleic Acids Research 50, D988–D995 (2022).

[25] Consortium, G. et al. The gtex consortium atlas of genetic regulatory effects across human tissues. Science 369, 1318–1330 (2020).

[26] Cohen, J. C., Boerwinkle, E., Mosley Jr, T. H. & Hobbs, H. H. Sequence variations in pcsk9, low ldl, and protection against coronary heart disease. New England Journal of Medicine 354, 1264–1272 (2006).

[27] Horton, J. D., Cohen, J. C. & Hobbs, H. H. Molecular biology of pcsk9: its role in ldl metabolism. Trends in Biochemical Sciences 32, 71–77 (2007).

[28] Dadu, R. T. & Ballantyne, C. M. Lipid lowering with pcsk9 inhibitors. Nature Reviews Cardiology 11, 563–575 (2014).

[29] Robinson, J. G. et al. Efficacy and safety of alirocumab in reducing lipids and cardiovascular events. New England Journal of Medicine 372, 1489–1499 (2015).

[30] Sabatine, M. S. et al. Evolocumab and clinical outcomes in patients with cardiovascular disease. New England Journal of Medicine 376, 1713–1722 (2017).

[31] Graham, S. E. et al. The power of genetic diversity in genome-wide association studies of lipids. Nature 600, 675–679 (2021).

[32] 1000 Genomes Project Consortium,. A global reference for human genetic variation. Nature 526, 68 (2015).

[33] Barton, A. R., Sherman, M. A., Mukamel, R. E. & Loh, P.-R. Whole-exome imputation within uk biobank powers rare coding variant association and fine-mapping analyses. Nature Genetics 53, 1260–1269 (2021).

[34] Kurki, M. I. et al. Finngen provides genetic insights from a well-phenotyped isolated population. Nature 613, 508–518 (2023).

[35] Hochner, H. et al. Parent-of-origin effects of the apob gene on adiposity in young adults. PLOS Genetics 11, e1005573 (2015).

[36] Ernst, J. & Kellis, M. Chromhmm: automating chromatin-state discovery and characterization. Nature Methods 9, 215–216 (2012).

[37] Ernst, J. & Kellis, M. Chromatin-state discovery and genome annotation with chromhmm. Nature Protocols 12, 2478–2492 (2017).

[38] Wachowski, N. A. et al. Implicating type 2 diabetes effector genes in relevant metabolic cellular models using promoter-focused capture-c. Diabetologia 1–14 (2024).

[39] Pautsch, A. et al. Crystal structure of glucokinase regulatory protein. Biochemistry 52, 3523–3531 (2013).

[40] Choi, J. M., Seo, M.-H., Kyeong, H.-H., Kim, E. & Kim, H.-S. Molecular basis for the role of glucokinase regulatory protein as the allosteric switch for glucokinase. Proceedings of the National Academy of Sciences 110, 10171–10176 (2013).

[41] Kichaev, G. et al. Leveraging polygenic functional enrichment to improve gwas power. The American Journal of Human Genetics 104, 65–75 (2019).

[42] Chen, M.-H. et al. Trans-ethnic and ancestry-specific blood-cell genetics in 746,667 individuals from 5 global populations. Cell 182, 1198–1213 (2020).

[43] Sakaue, S. et al. A cross-population atlas of genetic associations for 220 human phenotypes. Nature Genetics 53, 1415–1424 (2021).

[44] Garcia, C. K. et al. Autosomal recessive hypercholesterolemia caused by mutations in a putative ldl receptor adaptor protein. Science 292, 1394–1398 (2001).

[45] Wilund, K. R. et al. Molecular mechanisms of autosomal recessive hypercholesterolemia. Human Molecular Genetics 11, 3019–3030 (2002).

[46] Stayrook, K. R. et al. Regulation of carbohydrate metabolism by the farnesoid x receptor. Endocrinology 146, 984–991 (2005).

[47] Jiao, Y., Lu, Y. & Li, X.-y. Farnesoid x receptor: a master regulator of hepatic triglyceride and glucose homeostasis. Acta Pharmacologica Sinica 36, 44–50 (2015).

[48] Gomez-Ospina, N. et al. Mutations in the nuclear bile acid receptor fxr cause progressive familial intrahepatic cholestasis. Nature Communications 7, 10713 (2016).

[49] Green, R. C. et al. Acmg recommendations for reporting of incidental findings in clinical exome and genome sequencing. Genetics in Medicine 15, 565–574 (2013).

[50] Miller, D. T. et al. Acmg sf v3. 2 list for reporting of secondary findings in clinical exome and genome sequencing: a policy statement of the american college of medical genetics and genomics (acmg). Genetics in Medicine 25, 100866 (2023).

[51] Gao, B. & Zhou, X. Mesusie enables scalable and powerful multi-ancestry fine-mapping of causal variants in genome-wide association studies. Nature Genetics 56, 170–179 (2024).

[52] Zhang, Y., Qi, G.Park, J.-H. & Chatterjee, N. Estimation of complex effect-size distributions using summary-level statistics from genome-wide association studies across 32 complex traits. Nature Genetics 50, 1318–1326 (2018).

[53] Wang, L., Gao, B., Fan, Y., Xue, F. & Zhou, X. Mendelian randomization under the omnigenic architecture. Briefings in Bioinformatics 22, bbab322 (2021).

[54] Rolfe, E. D. L. et al. Association between birth weight and visceral fat in adults. The American Journal of Clinical Nutrition 92, 347–352 (2010).

