## Supplementary material for "Robust fine-mapping in the presence of linkage disequilibrium mismatch": RSparsePro_LD_note.pdf

#### 1 RSparsePro for robust fine-mapping

In RSparsePro, for a locus with $G$ variants, we assume the following data generating process:

$$
\begin{aligned}
\mathbf{s}_k &\sim \textbf{Multinomial}(1, \mathbf{1}_{G\times 1} \times \frac{1}{G}) \\
\beta_k &\sim \mathcal{N}(0, \sigma_\beta^2) \\
\mathbf{z} &\sim \mathcal{N}(\mathbf{R}\sum_k \mathbf{s}_k\beta_k, \mathbf{R}) \\
\hat{\mathbf{z}} &\sim \mathcal{N}(\mathbf{z}, \sigma_e^2\mathbf{I})
\end{aligned}
$$

where $\mathbf{s}_k$ is a $G \times 1$ vector serving as the sparse indicator for the causal variant in the $k^{th}$ credible set ($k \in \{1, \dots, K\}$); $\beta_k$ is the effect size of the $k^{th}$ credible set; $\mathbf{z}$ is the latent z-scores; and $\hat{\mathbf{z}}$ is the observed z-scores.

In robust fine-mapping, the objective is to infer the posterior probabilities of $\mathbf{s}_k$ based on the observed z-scores ($\hat{\mathbf{z}}$) and the linkage disequilibrium (LD) matrix ($\mathbf{R}$) derived from the reference panel. To achieve efficient and accurate inference, we follow our previous work [1, 2, 3] and develop a variational inference algorithm by approximating the desired posterior distribution with a paired mean field variational family [4, 5, 6].

We use a paired mean field factorized distribution

$$q(\mathbf{z}, \mathbf{s}_1, \ldots, \mathbf{s}_K, \beta_1, \ldots, \beta_K) = q(\mathbf{z}) \prod_k q(\mathbf{s}_k, \beta_k)$$

to approximate the desired posterior distribution

$$p(\mathbf{z}, \mathbf{s}_1, \ldots, \mathbf{s}_K, \beta_1, \ldots, \beta_K | \hat{\mathbf{z}}, \mathbf{R}) = \frac{p(\mathbf{z}, \mathbf{s}_1, \ldots, \mathbf{s}_K, \beta_1, \ldots, \beta_K, \hat{\mathbf{z}}, \mathbf{R})}{p(\hat{\mathbf{z}}, \mathbf{R})}$$

We can obtain the optimal approximation by maximizing the evidence lower bound (ELBO) [4]:

$$ELBO = E_{q(\mathbf{z}, \mathbf{s}_1, \ldots, \mathbf{s}_K, \beta_1, \ldots, \beta_K)}[\log \frac{p(\mathbf{z}, \mathbf{s}_1, \ldots, \mathbf{s}_K, \beta_1, \ldots, \beta_K, \hat{\mathbf{z}}, \mathbf{R})}{q(\mathbf{z}, \mathbf{s}_1, \ldots, \mathbf{s}_K, \beta_1, \ldots, \beta_K)}]$$

Specifically, to maximize the ELBO with respect to $\mathbf{z}$, we have [4]:

$$\begin{aligned}
\log q(\mathbf{z}) &= E_{q(\mathbf{s}_1, \ldots, \mathbf{s}_K, \beta_1, \ldots, \beta_K)}[\log p(\mathbf{z}, \mathbf{s}_1, \ldots, \mathbf{s}_K, \beta_1, \ldots, \beta_K, \hat{\mathbf{z}}, \mathbf{R})] \\
&= -\frac{(\mathbf{z} - \hat{\mathbf{z}})^2}{2\sigma_e^2} - \frac{(\mathbf{z} - \mathbf{R}\mathbf{h}^*)^T \mathbf{R}^{-1} (\mathbf{z} - \mathbf{R}\mathbf{h}^*)}{2} + const
\end{aligned}$$

where $\mathbf{h}^* = E_{q(\mathbf{s}_1, \ldots, \mathbf{s}_K, \beta_1, \ldots, \beta_K)}[\sum_k \mathbf{s}_k \beta_k]$. We can recognize that $q(\mathbf{z})$ follows a multivariate normal distribution

$$q(\mathbf{z}) \sim \mathcal{N}(\mathbf{u}_z^*, \mathbf{\Sigma}_z^*)$$

where

$$\mathbf{u}_z^* = \mathbf{R}(\frac{1}{\sigma_e^2}\mathbf{R} + \mathbf{I})^{-1}(\frac{1}{\sigma_e^2}\hat{\mathbf{z}} + \mathbf{h}^*)$$

$$\mathbf{\Sigma}_z^* = \mathbf{R}(\frac{1}{\sigma_e^2}\mathbf{R} + \mathbf{I})^{-1}$$

To maximize the ELBO with respect to $\mathbf{s}_k$ and $\beta_k$, similarly, we have:

$$\begin{aligned}
\log q(\mathbf{s}_k, \beta_k) &= E_{q(\mathbf{z}, \mathbf{s}_1, \ldots, \mathbf{s}_{k-1}, \mathbf{s}_{k+1}, \ldots, \mathbf{s}_K, \beta_1, \ldots, \beta_{k-1}, \beta_{k+1}, \ldots, \beta_K)}[\log p(\mathbf{z}, \mathbf{s}_1, \ldots, \mathbf{s}_K, \beta_1, \ldots, \beta_K, \hat{\mathbf{z}}, \mathbf{R})] \\
&= -\frac{(\beta_k)^2}{2\sigma_\beta^2} - \frac{(\mathbf{d}^* - \mathbf{R}\mathbf{s}_k\beta_k)^T \mathbf{R}^{-1} (\mathbf{d}^* - \mathbf{R}\mathbf{s}_k\beta_k)}{2} + const
\end{aligned}$$

where $\mathbf{d}^* = E_{q(\mathbf{z}, \mathbf{s}_1, \ldots, \mathbf{s}_{k-1}, \mathbf{s}_{k+1}, \mathbf{s}_K, \beta_1, \ldots, \beta_{k-1}, \beta_{k+1}, \ldots, \beta_K)}[\mathbf{z} - \mathbf{R}\sum_{k\prime \neq k} \mathbf{s}_{k\prime}\beta_{k\prime}]$.

We can recognize that

$$q(\beta_k | s_{k1} = 0, \ldots, s_{k(g-1)} = 0, s_{kg} = 1, s_{k(g+1)} = 0, \ldots, s_{kG} = 0) \sim \mathcal{N}(\mu_{kg}^*, \sigma_{kg}^{*2})$$

with

$$\mu^*_{kg} = \frac{d^*_g}{1 + \frac{1}{\sigma^2_\beta}}$$

$$\sigma^{*2}_{kg} = (1 + \frac{1}{\sigma^2_\beta})^{-1}$$

$$\gamma^*_{kg} := q(s_{k1} = 0, \ldots, s_{k(g-1)} = 0, s_{kg} = 1, s_{k(g+1)} = 0, \ldots, s_{kG} = 0) = softmax(\frac{\mu^{*2}_{kg}}{2\sigma^{*2}_{kg}})$$

With these iterative updates, we have an efficient variational algorithm for posterior inference summarized below in **Algorithm** 1.

**Algorithm 1:** RSparsePro for robust fine-mapping

```
Hyperparameters K, σ_β^2, σ_e^2
Data: ẑ, R
Result: γ*_kg, k ∈ {1, ..., K} and g ∈ {1, ..., G}
while not converge do
    for k = 1 to K do
        for g = 1 to G do
            update γ*_kg, μ*_kg;
        end
    end
    update u*_z;
end
```

#### 2 Hyperparameter settings

There are three hyperparameters in RSparsePro: $K$, $\sigma^2_\beta$ and $\sigma^2_e$. $K$ is the maximum number of credible sets in a locus. Similar to SuSiE [6], RSparsePro is robust to the choice of $K$, provided that it is greater than the actual number of causal variants. $\sigma^2_\beta$ is the hyperparameter for the prior distribution of the effect size. In practice, we set $\frac{1}{\sigma^2_\beta} = 0$, which corresponds to an uninformative prior. Notably, $\sigma^2_e$ is a critical hyperparameter, as it quantifies the discrepancy between the latent z-scores and the observed z-scores. If $\sigma^2_e$ is overly small and does not sufficiently capture the actual LD mismatch, the RSparsePro algorithm cannot converge. Conversely, if $\sigma^2_e$ is excessively large, RSparsePro may not adequately utilize the LD information, leading to reduced statistical power. Therefore, we first set $\sigma^2_e = 0$, with which the algorithm is equivalent to SparsePro [1]. If the algorithm does not converge, we then set $\sigma^2_e = 0.001$ and increase $\sigma^2_e$ by a factor of $1.5$ until the RSparsePro algorithm successfully converges.

### 3 Posterior summary

To construct 95% credible sets, we first calculate the attainable coverage ($AC$) of each credible set as in our previous work [1]:

$$AC_k = \sum_g \gamma_{kg}$$

where

$$\gamma_{kg} = \begin{cases} \gamma^*_{kg}, & \text{if } \gamma^*_{kg} = \max\left(\gamma^*_{1g}, \ldots, \gamma^*_{Kg}\right) \\ 0, & \text{otherwise} \end{cases}$$

Then, we add variants into the 95% credible set in descending order of the posterior probability $\gamma_{kg}$ for the $k^{th}$ credible set until $AC_k > 0.95$.

Additionally, for variant-level posterior inclusion probability (PIP), we have:

$$PIP_g = 1 - \Pi_k(1 - \gamma_{kg}) = 1 - (1 - \max(\gamma_{1g}, \ldots, \gamma_{Kg})) = \max\left(\gamma^*_{1g}, \ldots, \gamma^*_{Kg}\right)$$

for the $g$-th variant.

### References


[1] Zhang, W., Najafabadi, H. & Li, Y. Sparsepro: An efficient fine-mapping method integrating summary statistics and functional annotations. *PLOS Genetics* **19**, e1011104 (2023).

[2] Zhang, W., Sladek, R., Li, Y., Najafabadi, H. S. & Dupuis, J. Accounting for genetic effect heterogeneity in fine-mapping and improving power to detect gene-environment interactions with sharepro. *bioRxiv* 2023–07 (2023).

[3] Zhang, W. *et al.* Sharepro: an accurate and efficient genetic colocalization method accounting for multiple causal signals. *Bioinformatics* **40**, btae295 (2024).

[4] Blei, D. M., Kucukelbir, A. & McAuliffe, J. D. Variational inference: A review for statisticians. *Journal of the American Statistical Association* **112**, 859–877 (2017).

[5] Titsias, M. & Lazaro-Gredilla, M. Spike and slab variational inference for multi-task and multiple kernel learning. *Advances in Neural Information Processing Systems* **24**, 2339–2347 (2011).

[6] Wang, G., Sarkar, A., Carbonetto, P. & Stephens, M. A simple new approach to variable selection in regression, with application to genetic fine mapping. *Journal of the Royal Statistical Society Series B: Statistical Methodology* **82**, 1273–1300 (2020).
