## Supplementary figures and images for "Robust fine-mapping in the presence of linkage disequilibrium mismatch"

### SuppFigures.pptx

#### Slide 1
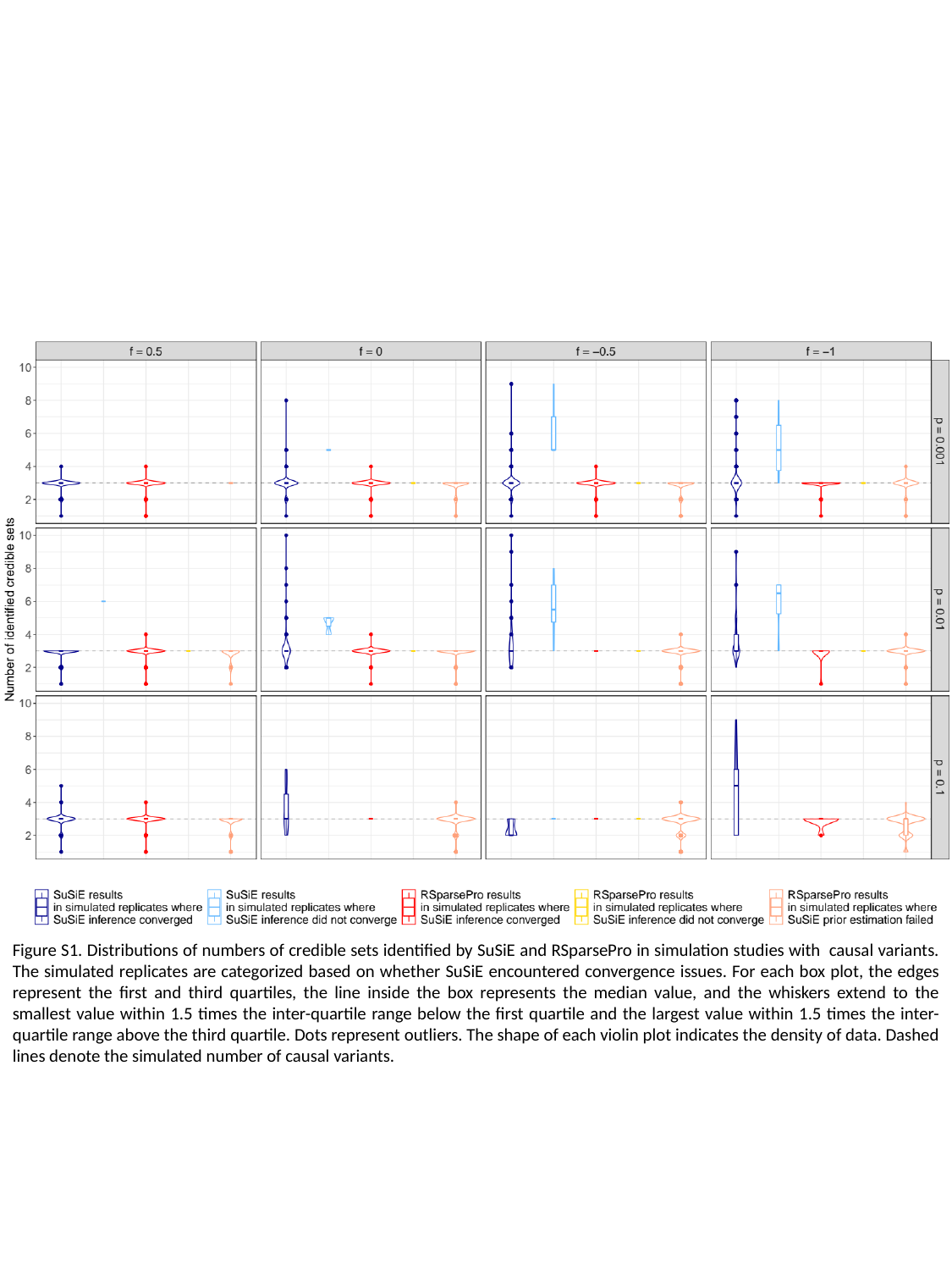

#### Slide 2
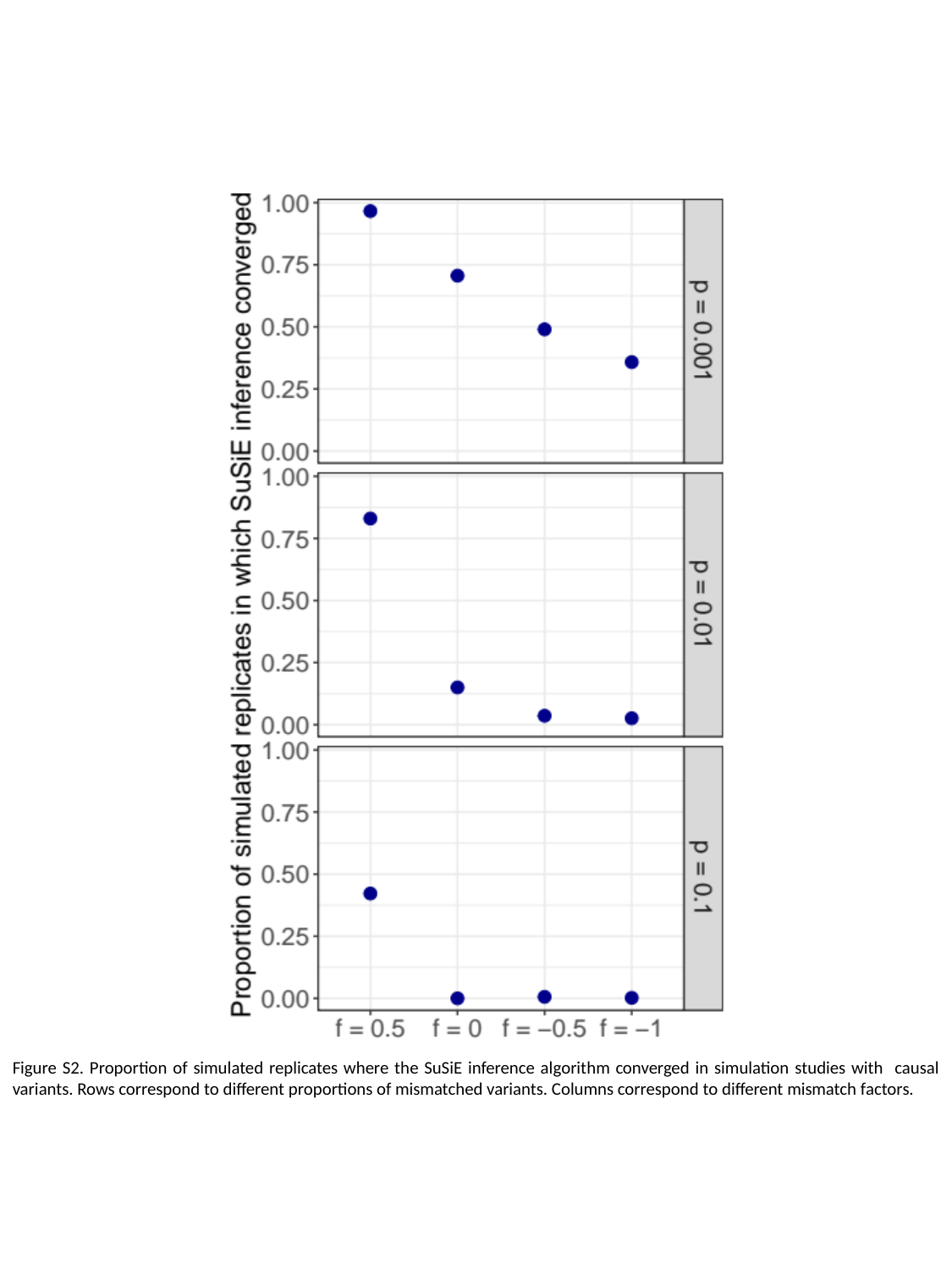

#### Slide 3
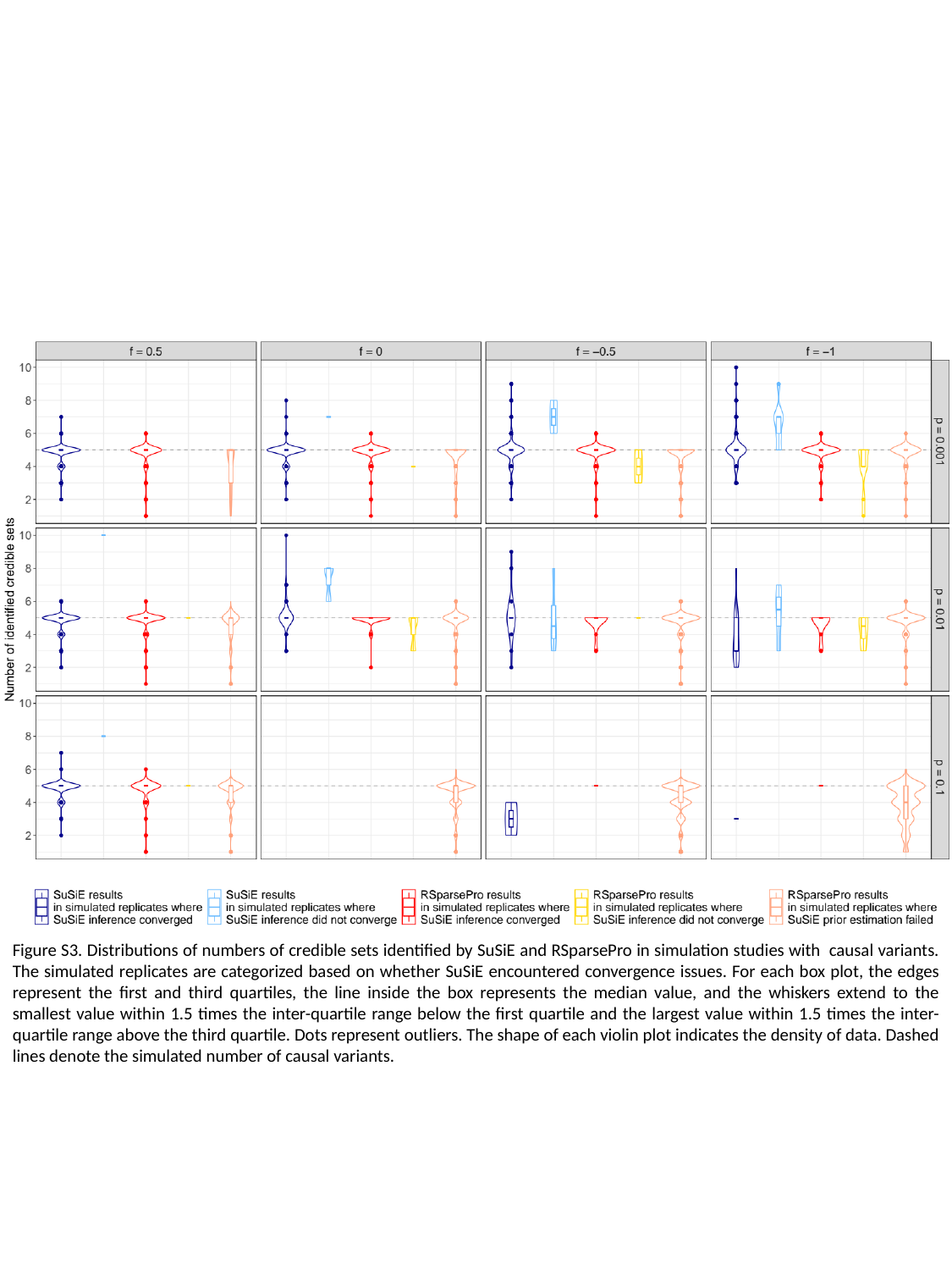

#### Slide 4
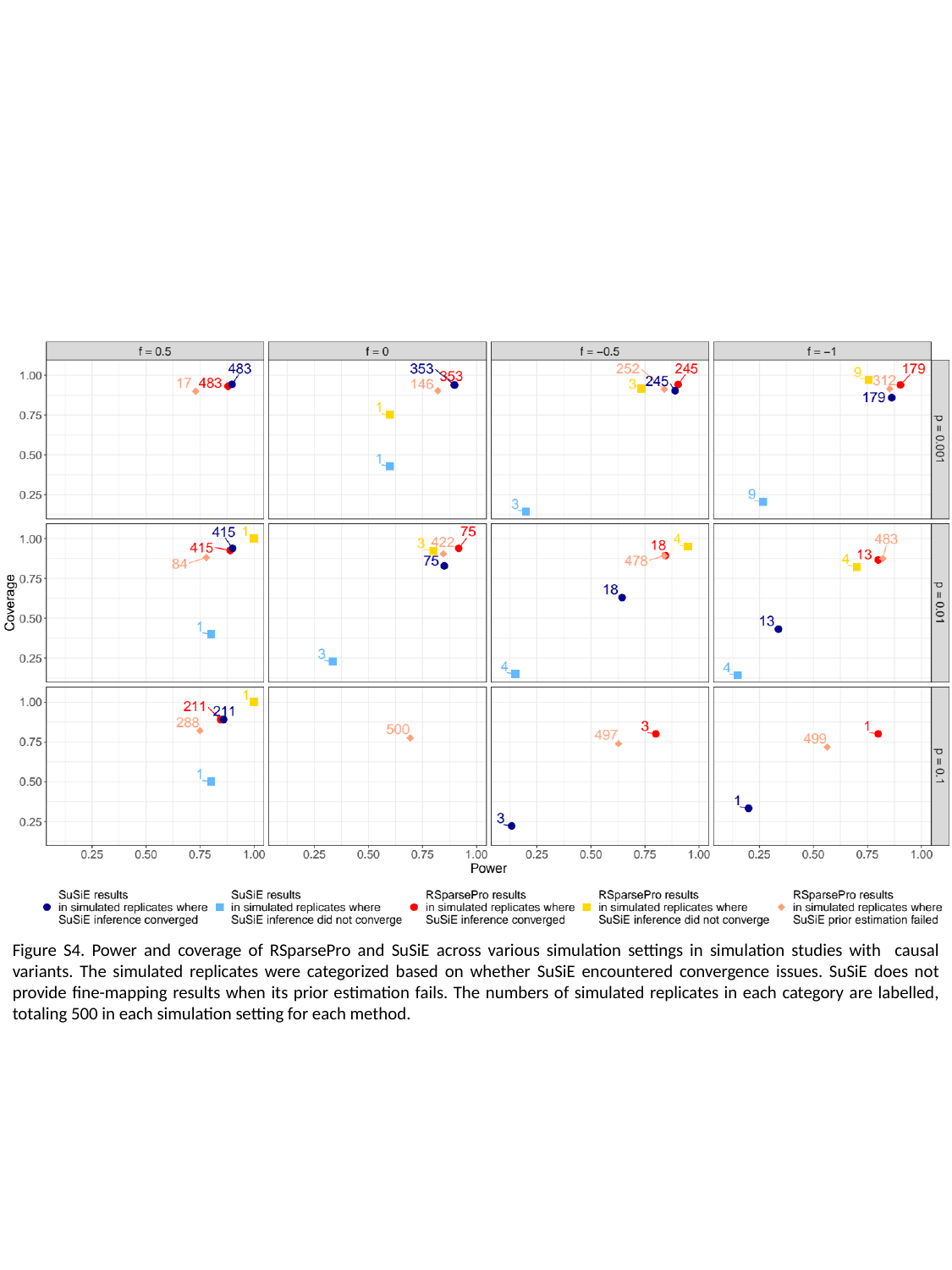
